# CRISPR/Cas9-mediated germline mutagenesis in the subsocial parasitoid wasp, *Sclerodermus guani*

**DOI:** 10.1101/2025.01.11.632282

**Authors:** Zi Ye, Guanzhen Fan, Yun Wei, Li Li, Feng Liu

## Abstract

The ectoparasitoid wasp *Sclerodermus guani* (Hymenoptera: Bethylidae), as a subsocial insect, is widely applied in biological control against beetle vectors of pine wood nematodes. Despite significant advances in behavioral research, functional genetics in *S. guani* remains underdeveloped due to the absence of efficient gene manipulation tools. In this study, we employed CRISPR-mediated mutagenesis to achieve germline gene knockout targeting the eye pigment associated gene *kynurenine 3-monooxygenase* (*KMO*). Phylogenetic analysis revealed that *S. guani* KMO shares a close relationship with its homolog in *Prorops nasuta* (Hymenoptera: Bethylidae). Two single-guide RNAs (sgRNAs), coupled with Cas9 protein with and without nuclear localization signal (NLS) were tested. Both sgRNAs induced specific *in vitro* DNA cleavage and *in vivo* heritable indels at the target genomic loci. Homozygous null mutant females and males exhibit a white-eye phenotype, which was identified during pupal stage. Optimal editing efficiency *in vivo* was achieved using the Cas9-NLS variant. Given the complication of germline gene editing in eusocial Hymenopterans, the application of CRISPR in the subsocial parasitoid wasp *S. guani* provides an accessible research platform for the molecular evolution of insect sociality.

## Introduction

Pine wilt disease is a deadly pine tree disease caused by the pine wood nematode *Bursaphelenchus xylophilus*, which is suggested native to North America and was introduced into Japan in the early 1900s, and subsequently spreading to other East Asian countries, including China and Korea in the 1980s (Zhao et al., 2008; Zhou et al., 2024). The nematodes induce damages to pine trees by feeding on their epithelial cells and are transmitted to healthy trees by the vector *Monochamus* beetles (Coleoptera: Cerambycidae), particularly the Japanese pine sawyer *M. alternatus* (Futai, 2013). Due to the wood-boring behavior of the beetles, biological control using parasitoid wasps has emerged as one of the most effective strategies for managing pine wilt disease (Wan et al., 2024; Wang et al., 2019).

*Sclerodermus guani* (Hymenoptera: Bethylidae) is a generalist ectoparasitoid wasp indigenous to China, widely adopted as biocontrol agent against *M. alternatus*. The female wasps detect their host via chemosensory cues and subsequently paralyze the host through venom injection (L. Li et al., 2015; Z. Li et al., 2015; Zhu, 2016). Following host paralysis, the wasps consume the host’s hemolymph to support oogenesis, after which the eggs are laid and undergo larval development, ultimately resulting in host mortality. The sex of *S. guani* is determined by haplodiploidy, where fertilized eggs develop into females and unfertilized eggs develop into males. In addition to their parasitic behaviors, *S. guani* exhibit subsocial characteristics, which distinguishes them from the majority of solitary parasitoid wasps (Tang et al., 2014). The adult females demonstrate various cooperative maternal behaviors that significantly enhance the survival rates of their offspring (Hu et al., 2012). Intraspecific aggression is also observed among females during competition for hosts (Guo et al., 2023). While numerous studies have investigated the behavioral biology of *Sclerodermus* wasps (Malabusini and Lupi, 2024), there remains a relative paucity in research regarding their molecular biology, largely due to the absence of efficient genetic manipulation tools.

To address this technical limitation, here we present a germline mutagenesis approach with CRISPR/Cas9 (Clustered regularly interspaced short palindromic repeats) in *S. guani* as a platform for future genetic studies. Hitherto, CRISPR/Cas9 has been successfully employed in several Hymenopteran insects (Chiu et al., 2020; Hu et al., 2019; Kohno et al., 2016; Konu et al., 2023; Yan et al., 2017), including solitary parasitoid wasps (Bai et al., 2024; Li et al., 2017a). However, limited studies have focused on inducing inheritable homozygous mutations in genes associated with social behaviors. Notably, in contrast to eusocial insects, which are typically challenging to genetically manipulate due to the complexities of establishing mutant lines (Kohno and Kubo, 2019), the subsocial nature of *Sclerodermus* wasps offers a promising model for investigating the molecular mechanisms of insect social interactions.

## Materials and Methods

### Insect rearing

The *S. guani* colony was provided by the Pest Control and Resource Utilization Laboratory at Guizhou Normal University (Wei et al., 2023). The parasitoid wasps were reared at 25°C, 65% humidity under a 12-hr light/12-hr dark photoperiod for more than 50 consecutive generations. Larvae of *M. alternatus* were purchased from Kaili, Guizhou province, China (107.981°E, 26.566°N), and were individually maintained in 5mL test tubes filled with sawdust at 4°C upon receiving.

### Bioinformatics

RNA extraction was performed on multiple 3–5-day old female adults that weigh approximately 0.3g in total using TRIzol Reagent (Invitrogen, 15596026) following the manufacturer’s protocol. The RNA sample was processed, validated, and sequenced by Biomarker Technologies Co., LTD. (Beijing, China) using an Illumina NovaSeq6000 sequencer (paired-end, 2 × 150 bp). The sequencing data were assembled *de novo* into transcripts following the Trinity protocol (Grabherr et al., 2011), and were searched against NR, Swiss-Prot, COG, KOG, eggNOG4.5, and KEGG databases for functional annotation.

The coding sequences of *KMO* genes were acquired from GenBank (https://www.ncbi.nlm.nih.gov/genbank/) and imported into MEGA11 (Tamura et al., 2021) for ClustalW multiple sequence alignment. Sequences from *Prorops nasuta* (XM_066724817.1), *Apis mellifera* (XM_026440256.1), *Nasonia vitripennis* (XM_001602208.5), *Drosophila melanogaster* (NM_078927.3), *Anopheles gambiae* (XM_315983.5), *Aedes aegypti* (XM_001653466.2), *Bombyx mori* (NM_001112665.1), *and Tribolium castaneum* (NM_001039411.1) were included in the analysis. The aligned protein sequences were analyzed to construct a phylogenetic tree using maximum likelihood with 1000 bootstrap replicates.

### sgRNA synthesis

As genomic information is currently unavailable for *S. guani*, the sgRNA sequences were designed using two different strategies. sgRNA1 was designed using the web tool CHOPCHOP (Labun et al., 2019) against the genome of the honeybee *A. mellifera*. The sgRNA1 sequence was selected based on its efficiency and sequence homology between *A. mellifera* and *S. guani*. sgRNA2 was designed using CRISPOR (Concordet and Haeussler, 2018) without genomic data. The sgRNAs were synthesized using EnGen sgRNA Synthesis Kit (NEB, E3322S) and purified with Monarch Spin RNA Cleanup Kit (NEB, T2030S) according to the manufacturer’s instruction (**Supplementary information 1**). sgRNA integrity and concentration were validated by gel electrophoresis and a microvolume spectrophotometer (Implen, N50).

### *In vitro* Cas9 cleavage assay

Commercially available Cas9 proteins were selected based on the presence or absence of nuclear localization signal (NLS): TrueCut Cas9 Protein (with NLS; Invitrogen, A36498) and *Spy* Cas9 Nuclease (without NLS; NEB, M0386T). Primers were designed to amplify the genomic regions flanking the sgRNA target sites via PCR (**Supplementary information 1**). The sgRNA and Cas9 protein were mixed at a molar ratio of 1.5:1 (320ng/µL Cas9 + 100ng/µL sgRNA), and incubated at room temperature for 20mins for ribonucleoprotein (RNP) assembly. 3µL PCR amplicons were incubated with 1µL RNP in a 30µL reaction at 37°C for 15mins. Negative controls were performed in parallel, omitting RNP from the reaction. DNA fragments were visualized by gel electrophoresis.

### Microinjection

Newly emerged *S. guani* adults were allowed to mate for 5-10 days. Mated females were collected to parasitize the larvae of *M. alternatus* at a ratio of 1 wasp per 0.1g host weight. At least 10 hosts were parasitized for each microinjection session. Female wasps generally require several days to paralyze their hosts and acquire nutrition for oogenesis by feeding on the host’s hemolymph. Once the female wasps begin lay eggs on the host cuticle, newly laid embryos were collected every hour and carefully aligned with fine forceps against the edge of a coverslip adhered to a microscope slide for injection.

The sgRNA and Cas9 protein were mixed at a molar ratio of 1.5:1 (320ng/µL Cas9 + 100ng/µL sgRNA or 160ng/µL Cas9 + 50ng/µL sgRNA) and incubated at room temperature for 20mins to assemble the RNP complex. The RNP was kept on ice and injected into the posterior end of the embryos (G0) using a FemtoJet 4i Microinjector (Eppendorf) and borosilicate needles (Sutter, BF100-50-10), which were prepared with a PC-100 Vertical Puller (Narishige). Injections were performed under a brightfield microscope equipped with a 10X objective lens (Sunny Optical, E5). The injected embryos were subsequently incubated on the slide under rearing conditions for 3 days.

*M. alternatus* larvae were freshly killed by freezing at -20°C for at least 3 days. The undamaged injected embryos were carefully placed on the cuticle of a dead host larva and further incubated under rearing conditions. Mature larvae (naturally dislodged from the hosts) were manually grouped together to simulate maternal care behaviors (Wu et al., 2017). The G0 virgin females were allowed to parasitize the hosts and reproduce through parthenogenesis. The resulting F1 male offspring were screened under a dissecting microscope (Sunny Optical, SZMN). White-eyed F1 males were collected and crossed with wild-type females to establish a homozygous mutant line.

### Microscopy

The images of *S. guani* developmental stages were captured using a dissecting microscope (Sunny Optical, SZMN) equipped with a digital camera.

### DNA extraction, PCR and sequencing

DNA extraction was performed either with TIANamp Genomic DNA Kit (TIANGEN, 4992254) following the manufacturer’s protocol, or a DNA extraction buffer (STE buffer: 10mg/mL proteinase K = 50:1). A pair of wings from a male or a single midleg from a female was gently removed and placed into 30µL DNA extraction buffer in a 0.5mL PCR tube. The sample was incubated in a thermocycler (Bio-Rad, C1000) at 65°C for 50mins and 95°C for 10mins. Extracted DNA samples were subjected to PCR and sequencing.

PCR was performed using 2x Rapid Taq Master Mix (Vazyme, P222-01) according to the standard protocol. PCR products were sequenced by Sangon Biotech (Shanghai, China).

## Results

### Kynurenine 3-monooxygenase sequence alignment and phylogenetic analysis

Kynurenine 3-monooxygenase (KMO), also known as kynurenine hydroxylase or cinnabar, is an enzyme that converts kynurenine to 3-hydroxy-kynurenine, a critical intermediate in the biosynthesis of the eye pigment ommochrome (Hebbar et al., 2023). *KMO* null mutants typically exhibit lighter eye color compared to the wild types due to the absence of brown pigment. *KMO* has been targeted in multiple insect species as an effective qualitative marker to assess gene editing efficiency (Feng et al., 2021; Li et al., 2017a, 2017b; Xu et al., 2020), and was therefore chosen for this study.

The full-length coding sequence of *KMO* gene in *S. guani* was obtained from an RNA-seq study (unpublished). The full-length coding sequence is 1,293bp that encodes 430 amino acids. Genomic DNA sequencing further reveal *S. guani KMO* gene consists of 10 exons (**Supplementary information 2**). The KMO amino acid sequence identity between *S. guani* and other selected species ranges from 50% to 87%, with the highest identity observed with *P. nasuta* (87%) (**Figure 1A**). Other species analyzed include *A. mellifera* (70%), *N. vitripennis* (67%), *D. melanogaster* (50%), *An. gambiae* (55%), *Ae. aegypti* (54%), *B. mori* (54%), and *T. castaneum* (50%). Phylogenetic analysis further confirms that the KMO protein in *S. guani* shares the closest evolutionary relationship with that in the Hymenopteran insect *P. nasuta* (**Figure 1B**), a solitary ectoparasitoid wasp from the Bethylidae family.

**Figure 1.**
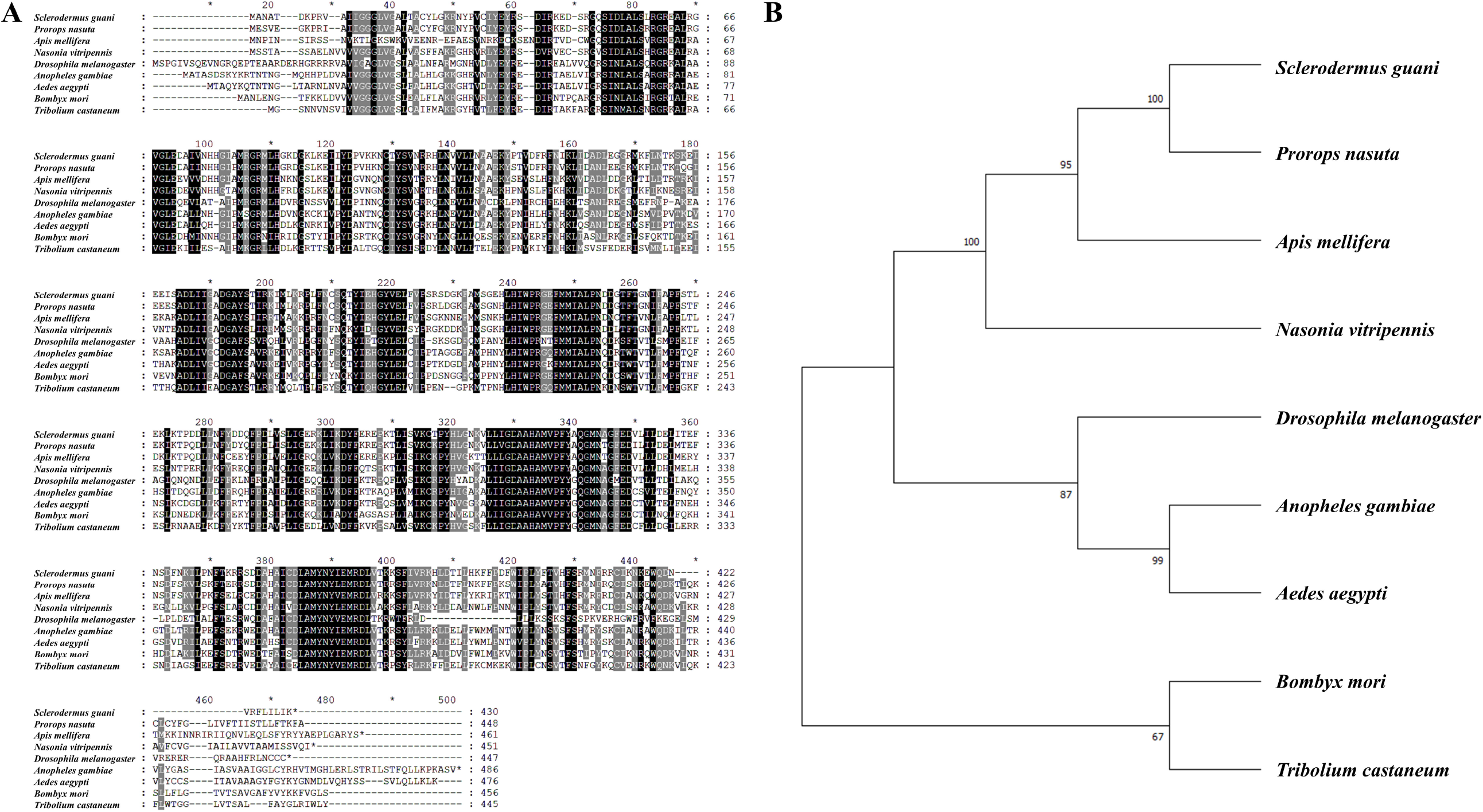
**(A)** Protein Sequence alignment of *S. guani* KMO with other insect species: Hymenoptera (*P. nasuta, A. mellifera, N. vitripennis*), Diptera (*D. melanogaster, An. gambiae, Ae. aegypti*), Lepidoptera (*B. mori*), and Coleoptera (*T. castaneum*). Residues in black indicate identical residues, and residues in grey indicate similar amino acid families in at least 80% of the aligned sequences. **(B)** Phylogenetic tree of KMO protein sequences from different species, inferred using Maximum Likelihood method. The percentage of replicate trees in which the associated taxa clustered together in the bootstrap test 1000 replicates are shown next to the branches.

### *In vitro* Cas9 cleavage validation

Based on the genomic structure of *KMO*, two sgRNAs were designed to target the 6th (sgRNA1) and the 3rd (sgRNA2) exons, respectively (**Figure 2A**). Previous studies have suggested that the presence of a nuclear localization signal (NLS) is crucial for the transport of Cas9 to the nucleus and is likely essential for efficient gene editing (Cong et al., 2013; Li et al., 2017a; Shirai et al., 2022). Therefore, both commercially available Cas9 proteins, one with NLS and one without, were tested in this study.

**Figure 2.**
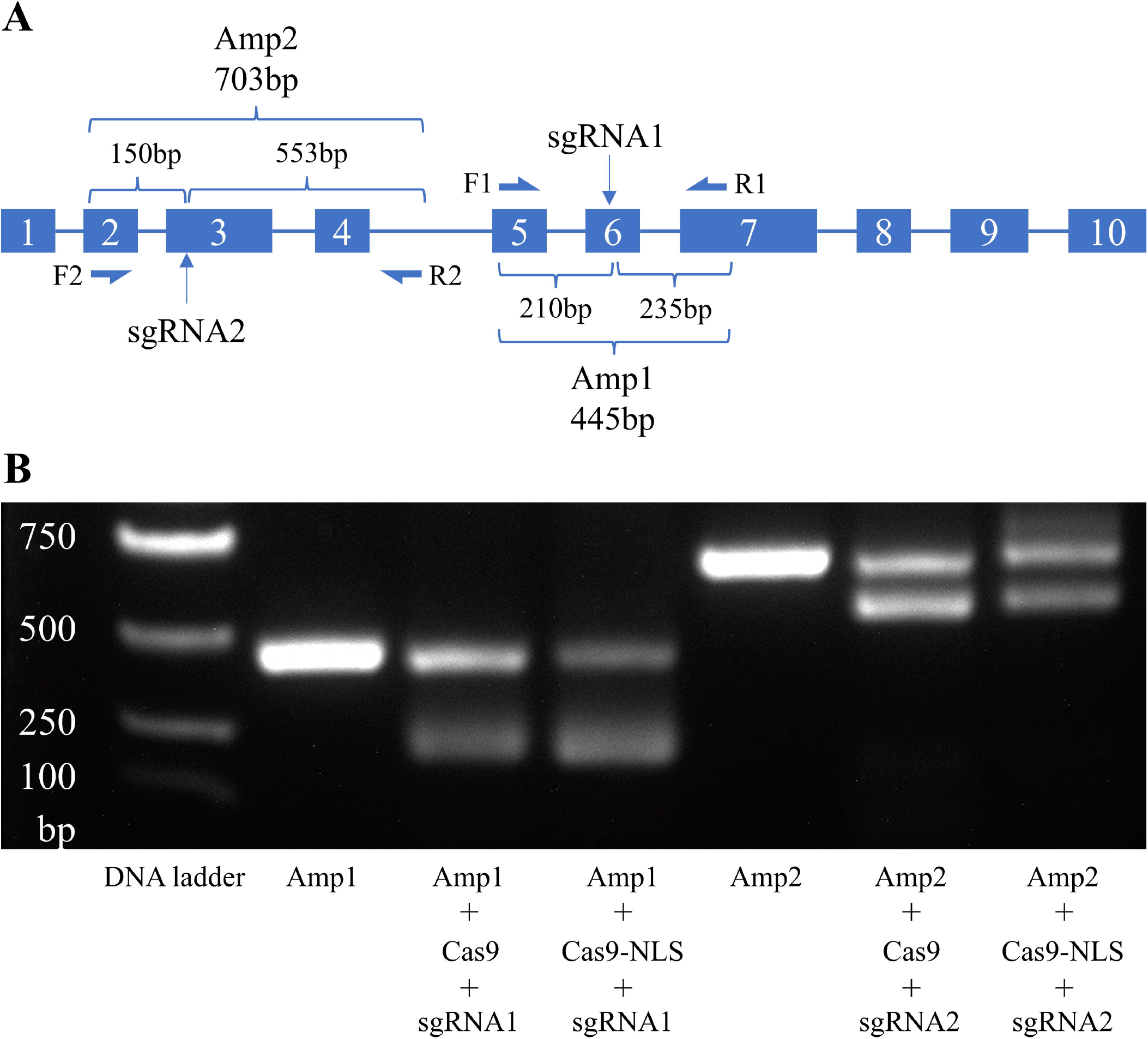
**(A)** Schematic of PCR strategy used to confirm mutagenesis. sgRNA1 and sgRNA2 were designed to target Exon 6 and Exon 3 of *KMO*, respectively. Two pairs of primers (*F1*/*R1* and *F2*/*R2*) were used to produce amplicons Amp1 (445bp) and Amp2 (703bp) via genomic PCR, respectively. **(B)** Agarose gel electrophoresis showing the amplicons after *in vitro* Cas9 cleavage assays.

To validate the activity and specificity of the sgRNAs and Cas9 proteins, we amplified the genomic regions flanking the target sites and assembled reactions *in vitro* to facilitate target cleavage. All sgRNA and Cas9 protein combinations induced effective target digestion, and the resulting DNA fragments were of the expected length (**Figure 2B**).

### Kynurenine 3-monooxygenase gene knockout

The pre-adult developmental stages of *S. guani* take approximately 3-4 weeks under our lab rearing conditions (Niu et al., 2023). The pupal stage is ideal for observing the development of compound eyes and ocelli, as eye pigment formation is not visible during earlier pre-adult stages (**Figure 3A-3I**). The posterior end of the egg (Figure 3A) corresponds to the abdomen of the larva (**Figure 3B**). *S. guani* females, being ectoparasitoids, lay their eggs on the cuticle of their hosts, which allows for easy collection and alignment of eggs for microinjection (**Figure 4A** and **4B**). The detailed embryonic injection protocol is described in Materials and Methods (**Figure 4C**).

**Figure 3.**
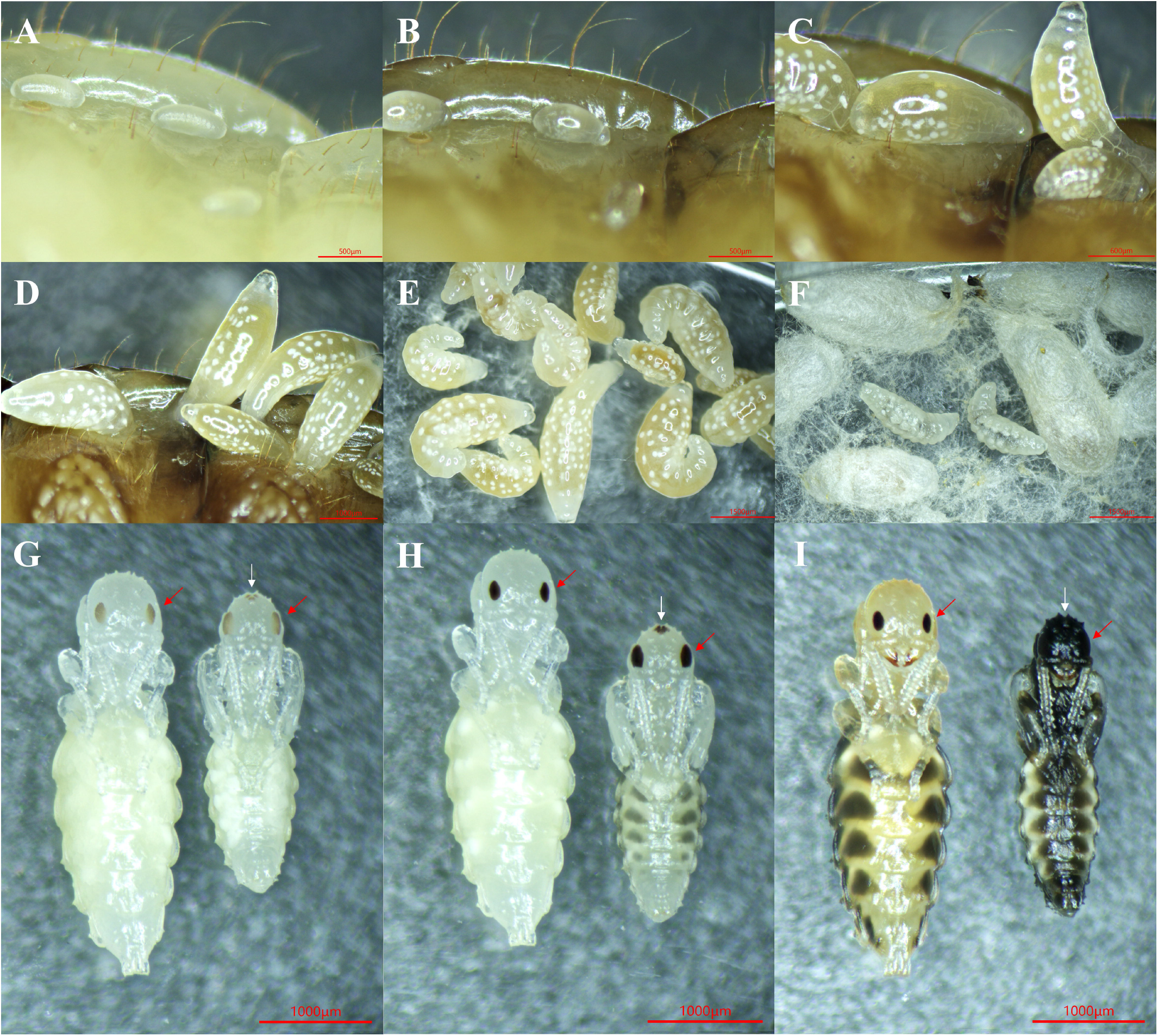
Representative developmental stages of pre-adult *S. guani*. Images were taken at following stages: egg **(A)**; larval stages on the 1st day **(B)**, 3rd day **(C)**, 5th day **(D)**, 7th day **(E)**, and 9th day **(F)**; pupal stages (left: female; right: male) on the 3rd day **(G)**, 7th day **(H)**, and 10th day **(I)**. Red arrows indicate the compound eyes and white arrows indicate the ocelli. Mature larvae were manually grouped together **(E)**.

**Figure 4.**
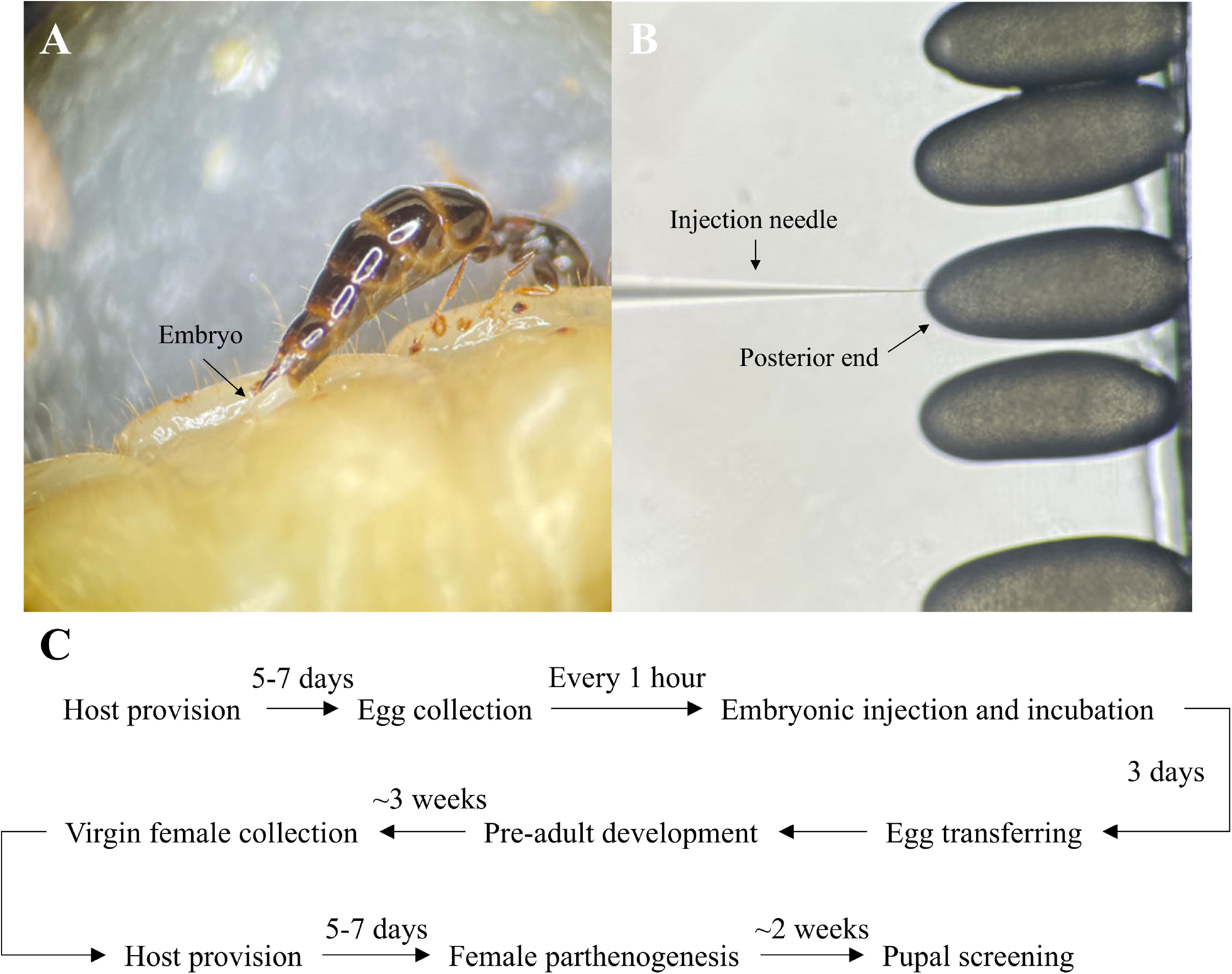
**(A)** Embryos were collected within 1 hour after oviposition by female wasps. **(B)** CRISPR reagents were injected to the posterior end of aligned embryos. **(C)** Schematic outlining the egg collection, microinjection, and mutant screening process in *S. guani*.

To test mutagenesis efficiency, two sgRNAs were complexed with Cas9 proteins with/without NLS. G0 females were allowed to reproduce parthenogenetically, resulting in all offspring being haploid males in the F1 generation. Both sgRNAs induced apparent mutant phenotypes in F1 males at the pupal stage (**Table 1**). The *KMO* null mutants failed to develop the brown eye pigment, instead exhibiting white compound eyes on approximately the 7th day of the pupal stage, compared to the wild types (**Figure 5A**). By the 10th day, the cuticle darkened and the contrast between the mutants and wild types become less distinct (**Figure 5B**). In male adults, the difference in eye color between the mutants and wild types was minimal (**Figure 5C**), suggesting that adults are not preferable for screening *KMO* null mutants. Genomic DNA PCR and sequencing from representative F1 males confirmed successful genetic modifications of the *KMO* gene, with nucleotide deletions observed compared to the wild-type genotype (**Figure 5D** and **5E**).

**Table 1.** Summary of the injection and mutagenesis mediated by CRISPR/Cas9 targeting *KMO* in *S. guani*.

| sgRNA | Cas9 | Concentration | Injected | Survivor (G0) | Mutant (F1) |
| --- | --- | --- | --- | --- | --- |
| sgRNA1 | SpCas9 | 320 ng/μL Cas9 +<br>100 ng/μL sgRNA | 50 | 7 females<br>4 males | 21/485 (4.3%) |
| sgRNA1 | SpCas9 | 160 ng/μL Cas9 +<br>80 ng/μL sgRNA | 50 | 14 females<br>3 males | 0/662 (0) |
| sgRNA1 | SpCas9-NLS | 320 ng/μL Cas9 +<br>100 ng/μL sgRNA | 50 | 2 females<br>1 male | 31/55 (56.4%) |
| sgRNA2 | SpCas9 | 320 ng/μL Cas9 +<br>100 ng/μL sgRNA | 50 | 5 females<br>1 male | 0/253 (0) |
| sgRNA2 | SpCas9 | 160 ng/μL Cas9 +<br>80 ng/μL sgRNA | 50 | 11 females<br>3 males | 0/637 (0) |
| sgRNA2 | SpCas9-NLS | 320 ng/μL Cas9 +<br>100 ng/μL sgRNA | 50 | 2 females | 15/44 (34.1%) |

**Figure 5.**
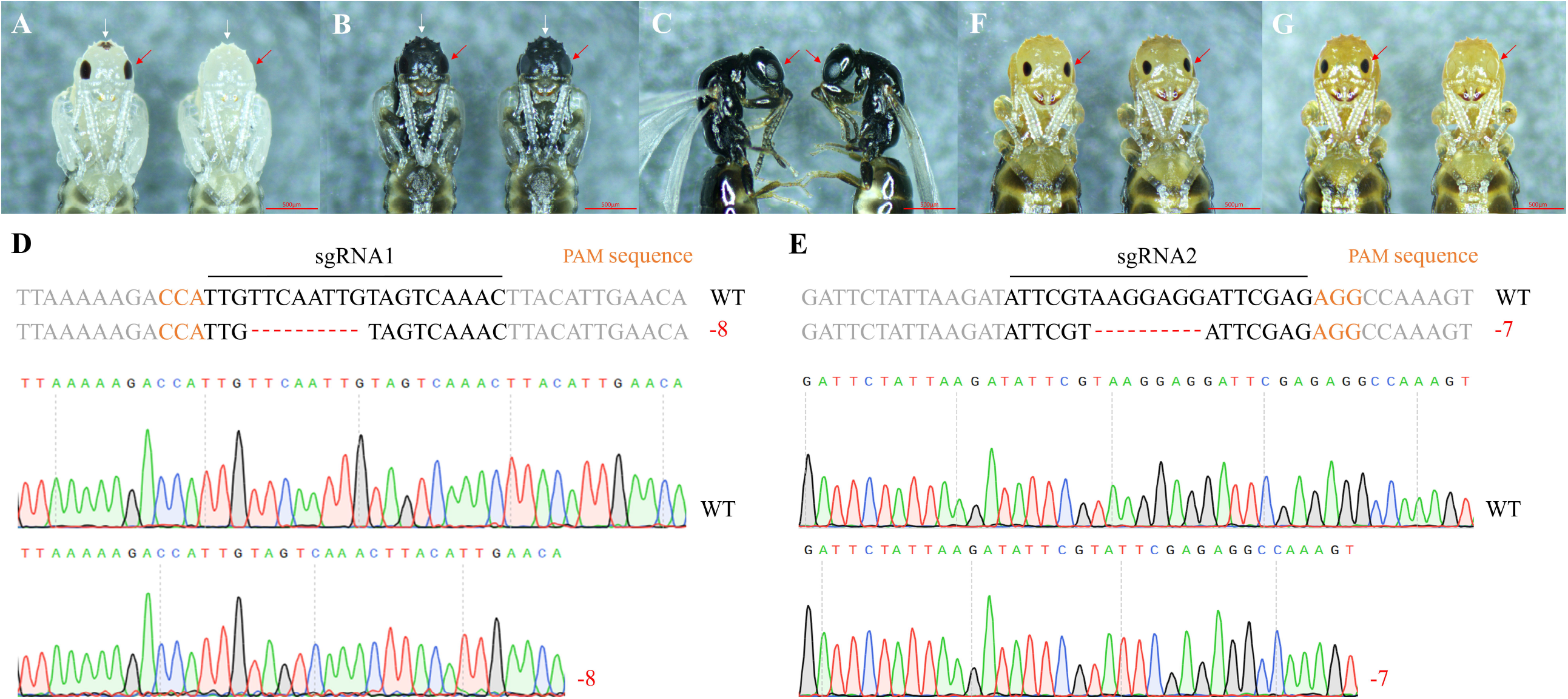
Phenotypes (left: wild type; right: *KMO* null mutant) of male pupae on the 7th day **(A)** and 10th day **(B)** of the pupal stage, and in adult males **(C)**. Sanger sequencing results confirm the -8 and -7 nucleotide deletions induced by sgRNA1 **(D)** and sgRNA2 **(E)**, respectively. sgRNA sequences are indicated in black, PAM sequences (NGG) are shown in orange, and indels are marked with red dashes. Phenotypes (left: wild type; right: *KMO* null mutant) of F2 heterozygous **(F)** and F3 homozygous **(G)** female pupae on the 10th day of the pupal stage. Red arrows indicate the compound eyes and white arrows indicate the ocelli.

To optimize mutagenesis efficiency, various reagent combinations were tested. We noticed that the most time-consuming step was egg collection, as the eggs are deposited singly. Consequently, injecting 50 embryos per session balanced efficacy with effort. sgRNA1 at 100 ng/µL was capable of generating mutants when complexed with either 320 ng/µL Cas9 or Cas9-NLS, while sgRNA2 at 100 ng/µL was only effective when complexed with 320 ng/µL Cas9-NLS. Compared to Cas9, Cas9-NLS exhibited higher mutagenesis efficiency but resulted in fewer G0 survivors. Both sgRNAs failed to generate mutants at 50 ng/µL when complexed with 160 ng/µL Cas9 (**Table 1**).

In order to further confirm the heritability of the mutations, F1 white-eyed males with indels induced by sgRNA1 were crossed with wild-type females. The resulting F2 heterozygous female pupae exhibited black compound eyes, which is comparable to the wild types (**Figure 5F**). Ten F2 females were randomly selected and sequenced to confirm their mutations without compromising their reproduction capability. The heterozygous female mutants were then crossed with white-eyed mutant males to generate homozygous females. The resulting F3 homozygous female pupae displayed expected white eye phenotype (**Figure 5G**), indicating that the mutation is recessively inherited in subsequent generations (**Table 2**).

**Table 2.** Summary of offspring phenotypes in F2 and F3 generations. F2 wasps were produced by crossing F1 white-eyed males with wild-type females. F3 wasps were generated by crossing F2 heterozygous females with white-eyed males.

| Phenotype | F2 female | F2 male | F3 female | F3 male |
| --- | --- | --- | --- | --- |
| +/+ (+ for male) | 0 | 15 (100%) | 0 | 7 (53.8%) |
| +/- | 10 (100%) | - | 89 (54.3%) | - |
| -/- (- for male) | 0 | 0 | 75 (45.7%) | 6 (46.2%) |

## Discussion

The CRISPR/Cas9 system has rapidly become one of the most versatile and efficient tools in insect genetic modification due to its amenability and ease of application. Although it facilitated germline mutagenesis across a wide range of insect taxa, its application in eusocial Hymenopterans remains challenging due to their unique reproductive characteristics (Singh and Dhar, 2020); that is, only the reproductive castes (queen and males) contribute to colony propagation, making the mutagenesis of sterile workers typically somatic and nonheritable. The first report of heritable mutagenesis in Hymenoptera using CRISPR/Cas9 was published in 2017, where the *cinnabar* gene was knocked out in the jewel wasp *N. vitripennis* (Li et al., 2017a). The same year, a significant advance was made in the Indian jumping ant *Harpegnathos saltator* (Yan et al., 2017). The researchers exploited the unique “pseudoqueen” mechanism of the species, allowing the mutant workers to become reproductive. While CRISPR/Cas9 has since been applied to other solitary ectoparasitoid wasps, such as *Habrobracon hebetor* (Bai et al., 2024), it remains inapplicable for generating germline mutations in eusocial Hymenopterans like honeybees, which possess a typical caste system.

In this study, we have developed a protocol for inducing programmable mutagenesis in the subsocial ectoparasitoid wasp *S. guani*. This advance opens up new avenues for investigating the molecular mechanisms underlying social behaviors in the parasitoid wasp. Compared to previous methods that require the injection of hundreds of embryos (Bai et al., 2024; Li et al., 2017b), our protocol demonstrates that heritable mutations can be generated with as few as 50 injected embryos, significantly reducing the experimental effort required. Furthermore, our results show that Cas9-NLS is more efficient than wild-type Cas9 in inducing mutations, which suggests that the NLS-tagged Cas9 may be more suitable for future applications. Although the survival rate of G0 individuals is lower when using Cas9-NLS, a finding consistent with other studies (Hu et al., 2018), increasing the number of embryos injected would likely compensate for this issue and yield a sufficient number of mutant founders.

The CRISPR/Cas9 system induces blunt double-stranded break (DSB), which is subsequently repaired through one of two endogenous DNA repair pathways: non-homologous end joining (NHEJ) or homology-directed repair (HDR) (Doudna and Charpentier, 2014). With the introduction of exogenous homology templates, specific DNA sequences can be knocked into the targeted genomic loci. Depending on the experimental purpose, various sizes of DNA fragments can be integrated. For instance, large genetic elements, such as fluorescence markers and transcription factors, can be carried by plasmid DNA templates, while point mutations can be introduced via single-stranded oligodeoxynucleotides (Gantz and Akbari, 2018). In the context of using parasitoid wasps as biological control agents, CRISPR-mediated HDR offers a promising strategy for strain optimization by integrating beneficial genetic regulators that enhance fitness or parasitic efficiency. The accessibility of HDR in *S. guani* could be explored by including HDR template in the microinjection complex to insert tissue-specific fluorescence markers, contingent on the availability of genomic data (Gratz et al., 2014). Moreover, our protocol could enable investigations into other transgenesis systems in *S. guani*, such as *piggyBac* and *phiC31* (Huang et al., 2016), thereby expanding the molecular toolkit for both research and biological control practices.

Embryonic injection remains the most commonly used method for delivering CRISPR machineries into insects to induce germline mutagenesis. Recently, several adult-injecting techniques were developed for oocyte delivery, such as Direct Parental CRISPR (DIPA) and Receptor-Mediated Ovary Transduction of Cargo (ReMOT) (Chaverra-Rodriguez et al., 2020, 2018; Shirai et al., 2022). In these techniques, CRISPR reagents are injected into females during vitellogenesis, where they are taken up by oocytes through endocytosis. These methods require the identification of female undergoing vitellogenesis, which is likely feasible in *S. guani*. Once the females have paralyzed their host, they spend several days feeding on the host’s hemolymph to support oogenesis, during which their abdomen visibly enlarged. This typical oogenesis process could potentially make adult injection techniques applicable in laboratories that lack advanced microinjection equipment.

## Author contributions

Z.Y., L.L., and F.L. designed research; Z.Y., G.F., and Y.W. performed research; Z.Y., G.F., Y.W., L.L., and F.L. analyzed data; Z.Y., G.F., Y.W., L.L., and F.L. wrote the paper.

## Acknowledgments

We thank all members of the F.L. and L.L. lab for suggestions and comments. We thank Guizhou Normal University and Shenzhen Bay Laboratory for equipment supports. This work was funded by National Natural Science Foundation of China (Grant No. 32401587), Guizhou Science and Technology Department (Grant No. Qian Ke He Ji Chu ZK [2024] General 423), Guizhou Education Department (Grant No. Qian Jiao Ji [2024] 54), and Guizhou Normal University (Grant No. QSXM [2022] B09).

## Disclosure

The authors declare no conflicts of interest.

## Notes

### Competing Interest Statement

The authors have declared no competing interest.

