## Supplementary information 2 for "CRISPR/Cas9-mediated germline mutagenesis in the subsocial parasitoid wasp, *Sclerodermus guani*"

Exons

Introns

>SG_KMO

ATGGCGAATGCAACGGATAAACCTCGAGTAGCTATTATCGGGGGTGGTTTAGTATGTTTCTCATTATTTT
AAACAATTCACACACACCCTAATTTGTTCGGATTTTTTTCTTACATTATTTATTGATTTTTTGTCAGGTTG
GAGCTTTAACAGCTTGTTATTTGGGCAAGCGGAATTATCCTGTGTGTATTTACGAATATCGTTCAGGTAA
AACATTCGATGTAACTTGTTGTAAACGTTCTTCCGAATTGCAATGACTCCATCAAGTATTTTGATATAATG
ATTCTATTAAGATATTCGTAAGGAGGATTCGAGAGGCCAAAGTATAGATCTTGCGCTATCTCTCCGTGGT
CGAGAAGCTCTTCGCGGGGTAGGTCTTGAAGATGCAATTGTTAATCATCACGGAATCGCCATGAGAGG
CAGGATGCTACATGGAAAAGATGGTAAACTCAAGGAAATTATTTATGACCCAGTAAAGAAAAATGTAA
GTACTAGAATTTCATTTATCAAAATGAAGAATATTTTATCGTTTCCCTCTCACTGTCATTTGCAATTTGCG
TATCATATACTTCTTAATCTTGTTTGGCAGTGCACATATTCTGTTAATCGAAGGCATCTTAATGTGGTTTTA
TTGAATGGTAAGTGTAGAGAAATGATTTCATTTAATTAAAAGTTCGTATTTCGAGGTGAGATGGCGAGA
GAAAGAGATTACGGGTGCATGTTTGTTAATTGATTGCTTATGTAATTAGTGATCTTGATATTTATGCACGA
AATCCTATAGGGCTGGCGGTAAAAGTGAGATACTTTATAATCTAAAAAAGTAGTCAATATCCTATTACGA
GAAGGTGCGTGAAGCCACAGAAATACCTGTGGATTATATTACACCCTTAGGTATTTAATAAAGAAAAAA
CAGTTTATTATTCAAAAAGATCAGAAATTTTGAAATTCTTTCATCATTCTTAATCATGACTAAATGGGTAT
CCATTTGATAGAAGAGAGTCGATATTATCATAGAAAAAACTGTTTCATGTTCTTTATTGTTCTTTGCTAAT
AGCTGCAGAAAAATACCCTACGGTGGATTTTCGTTTTAACATTAAATTAATCGACGCAGATCTTGAAGG
TGGAAGAATGAAATTTCTCAAGTAAGAAAATATAACAAATTTGTGTCTGACCTTTATTATTAATAATTGG
AATTTTAATTTTATTGAATTTTAGTACAAAGAGTAAGGAAATAGAGGAGATTTCTGCAGATTTGATTATT
GGAGCAGACGGAGCTTATTCGACAATACGGAAAATAATGTTAAAAAGACCATTGTTCAATTGTAGTCAA
ACTTACATTGAACACGGTTATGTAGAATTGTTTGTTCCGTCCAGATCCGATGGAAAAGTAATATAATGAT
TGTAACAAATTTAATATGCTTTAAACAAGAATATTTCAATCAGTCAAATTTATAATTTTAGTTTGCAATGA
GTGGGGAACATTTACATATTTGGCCTCGTGGTGAATTCATGATGATTGCTTTACCAAATGATGATGGCAC
GTTCACAGGAAATATCTTTGCACCCTTTTCCACTCTCGAGAAGTTGAAAACCCCCGATGATTTGCTTAAT
TTCTATGATGATCAATTTCCTGATTTGGTATCATTAATTGGAGAACGAAAGTTGATTAAGGATTATTTTGA
GAGGGAACCTAAAACTTTAATTTCTGTAAAGGTATAAAGTTCTTCGGTATAATATTCTTTTATTTTTTTAG
TCTTATTTTTTTTTTTTTAAATTAATTATATTATAATATATCAGTGCACTCCTTATCATCTTGGAAACAAGGT
TCTTCTTATTGGTGACGCTGCCCATGCAATGGTACCATTTTATGCCCAAGGAATGAATGCCGTAAATTTG
AATAATTAAATCATTATTAAATGATGAATATTCGTAGAAATACATTTTAATAATTAATTTTTTAAATTATTTG
TTTAATAGGGTTTCGAAGACGTTTTAATTTTGGATGAATTGATAACTGAATTTAATTCAGATTTCAACAA
AATTTTACCAAACTTTACAAAGCGTAGGTCAGACGATGCTCACGCAATTTGTGACCTTGCAATGTATAAT
TATATAGAGGTGAATTAGTTGAGTAAAATTAATTTCATTTGTATAACGTGTAACTAAATCTTTGTATGAAA
CATTGAATTTTAGATGAGAGACCTTGTAACAAAAAAGTCATTCATCGTAAGAAAACATTTGGATACGAT
TTTACACAAATTCTTCCCAGACTTCTGGATTCCTCTTTATTTTACAGTACACTTTTCAAGAATGAATTTTC
GACGGTGCATAAAAAATAAAGAATGGCAGGATAATGTAAGATTTCTAATTTTAATAAAATAA
